# G-OnRamp: A Galaxy-based platform for creating genome browsers for collaborative genome annotation

**DOI:** 10.1101/499558

**Authors:** Yating Liu, Luke Sargent, Wilson Leung, Sarah C.R. Elgin, Jeremy Goecks

**Affiliations:** Washington University in St. Louis, St. Louis MO 63130, USA; Oregon Health & Science University, Portland, OR 97239, USA

## Abstract

**Summary:** G-OnRamp provides a user-friendly, web-based platform for collaborative, end-to-end annotation of eukaryotic genomes using UCSC Assembly Hubs and JBrowse/Apollo genome browsers with evidence tracks derived from sequence alignments, *ab initio* gene predictors, RNA-Seq data, and repeat finders. Researchers can use the G-OnRamp output to visualize large genomics datasets, and can utilize the output to drive collaborative genome annotation projects in both research and educational settings.

**Availability and Implementation:** The G-OnRamp virtual machine images and tutorials are freely available on the G-OnRamp web site (http://g-onramp.org/deployments). The G-OnRamp source code is freely available under an Academic Free License version 3.0 through the goeckslab GitHub repository (https://github.com/goeckslab).

**Supplementary information:**

## 1. Introduction

As the cost continues to decrease, many more eukaryotic genomes are being sequenced (Cheng *et al*., 2018; Koepfli *et al*., 2015; i5K Consortium, 2013). The first task after genome sequencing is often annotating the genome with the locations of functional elements such as genes (exons, introns, splice sites) and promoters. High quality genome annotations are critical for understanding organism function and evolution. Genome browsers facilitate annotation, enabling researchers to visually synthesize results from multiple experiments. Genome annotation requires the integration of multiple lines of evidence, both experimental (*e.g*., RNA-Seq) and computational (*e.g*., gene predictions, sequence alignments) (Yandell and Ence, 2012). However, generating the evidence tracks and visualizations needed to analyze eukaryotic genomes remains technically challenging and time consuming.

Many analysis workflows for genome annotation are already available (Haas *et al*., 2008; Hoff *et al*., 2016; Holt and Yandell, 2011), but these workflows are often difficult for biologists with limited bioinformatics expertise to use. Key challenges include configuring needed tools and software dependencies, learning to use command-line tools, running tools on a computing cluster, converting the results for visualization [*e.g*., using the UCSC Genome Browser (Kent *et al*., 2002) or JBrowse (Buels *et al*., 2016)], maintaining web servers with the visualization platform to facilitate collaboration with other researchers, and keeping track of analysis steps and program parameters in order to re-use the workflow on other genome assemblies.

To address these challenges, the Genomics Education Partnership (GEP; http://gep.wustl.edu) has collaborated with Galaxy (https://galaxyproject.org/) to develop G-OnRamp, a scalable user-friendly web-based platform for eukaryotic genome annotation. G-OnRamp has computational workflows that combine more than 25 community and custom analysis tools to create evidence tracks leading to complete UCSC Assembly Hubs (Raney *et al*., 2014) and JBrowse/Apollo genome browsers. The output produced by G-OnRamp includes evidence tracks for homologous protein and transcript sequence alignments, *ab initio* gene predictions, transcriptional activity (based on RNA-Seq) with full transcripts and splice junctions, and repeats. G-OnRamp can be deployed on the Amazon Elastic Compute Cloud (EC2) via CloudLaunch (Afgan, Lonie, *et al*., 2018), or locally via a virtual appliance. Genome browsers created with G-OnRamp can be exported to the CyVerse Data Store (Merchant *et al*., 2016), where they can be used for annotation without the need to maintain a web server. G-OnRamp can convert a JBrowse genome browser into an Apollo instance (Lee *et al*., 2013) to enable real-time, collaborative genome annotations. Training materials and documentations for G-OnRamp are available on the G-OnRamp web site (http://g-onramp.org/training).

## 2. Features

Galaxy is an open, web-based platform for accessible, reproducible, and transparent analyses of large biological datasets that is used by thousands of scientists throughout the world (Afgan, Baker, *et al*., 2018). G-OnRamp extends Galaxy by providing the data analyses and conversions needed for constructing genome browsers for annotation.

G-OnRamp encapsulates the steps required to construct UCSC Assembly Hubs and JBrowse genome browsers into Galaxy workflows. Users need only to specify the input datasets—a genome assembly of the target genome, transcript and protein sequences from a closely-related informant genome, and RNA-Seq data from the target genome—and then run the workflow to create the genome browser. G-OnRamp can be used to generate a genome browser for annotation of almost any eukaryotic genome.

### 2.1 Primary tools in each sub-workflow

The G-OnRamp workflow consists of four sub-workflows: homologous sequence alignments, *ab initio* gene predictions, RNA-Seq analysis, and repeat identification. Each sub-workflow is composed of multiple bioinformatics tools. The outputs from these sub-workflows provide the input data for novel Galaxy tools, the Hub Archive Creator and the JBrowse Archive Creator, which construct the UCSC Assembly Hub and the JBrowse genome browser for the target genome (Figure 1). The key components of each sub-workflow are described below; more details are available at http://g-onramp.org.

**Figure 1:**
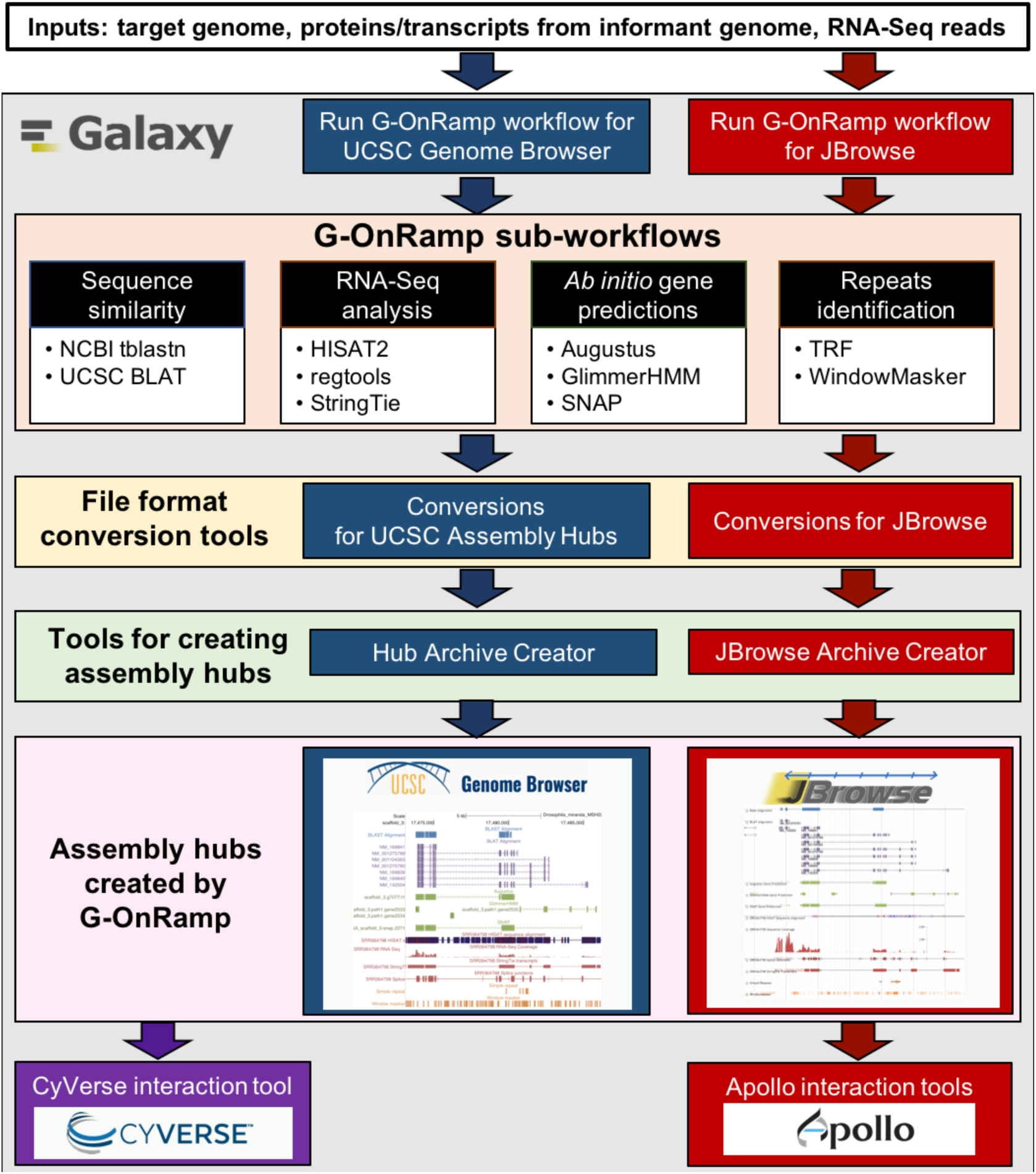
Overview of the G-OnRamp workflows. A researcher provides the target genome assembly for annotation, transcript and protein sequences from an informant genome, and RNA-Seq data from the target organism as input datasets to G-OnRamp. These datasets are processed by the sequence similarity, RNA-Seq analysis, *ab initio* gene predictions, and repeats identification sub-workflows to create UCSC Assembly Hubs and JBrowse genome browsers. The Apollo interaction tools can convert the JBrowse genome browser into an Apollo instance for collaborative annotation. The CyVerse interaction tool can transfer the G-OnRamp output to the CyVerse Data Store for long-term storage and visualization.

#### Homologous sequence similarity

This sub-workflow uses NCBI tblastn (Gertz *et al*., 2006; Camacho *et al*., 2009) and BLAT (Kent, 2002) to align proteins and transcripts from an informant genome to the target genome. BLAT transcript alignments are filtered to determine the locations of the putative orthologs. The “UCSC Trix Index Generator” tool creates an index that enables users to search for protein and transcript matches by name.

#### RNA-Seq analysis

This sub-workflow uses HISAT2 (Kim *et al*., 2015) to align RNA-Seq reads to the target genome. RNA-Seq read alignments are aggregated by the “Convert BAM to BigWig” tool to create a read coverage track. The “junctions extract” subprogram in regtools (https://github.com/griffithlab/regtools) is used to identify putative splice junctions. StringTie (Pertea *et al*., 2015) is used to construct putative transcripts from the aligned RNA-Seq reads.

#### *Ab initio* gene predictions

This sub-workflow consists of three gene predictors: Augustus (Stanke *et al*., 2006), GlimmerHMM (Majoros *et al*., 2004), and SNAP (Korf, 2004). Species-specific gene prediction parameters can be specified when the workflow is run.

#### Repeats identification

Tandem Repeats Finder (Benson, 1999) and the TrfBig utility developed by the UCSC Genome Bioinformatics Group are used to identify tandem repeats. WindowMasker (Morgulis *et al*., 2006) is used to identify transposon remnants and simple repeats.

### 2.2 Tools for constructing genome browsers

We have developed custom Galaxy tools, the Hub Archive Creator (HAC) and the JBrowse Archive Creator (JAC), that organize and format output data from the annotation sub-workflows so it is suitable for constructing new UCSC and JBrowse genome browsers. Users can customize names and colors of each evidence track, and assign related evidence tracks to the same group. Users can also incorporate custom evidence tracks (in GFF3, BED, GTF, BAM, and BigWig formats) into the genome browsers.

### 2.3 Collaborative annotation with Apollo

The “Create or Update Organism” tool creates an Apollo instance from the JAC output. The “Apollo User Manager” tool provides batch management by administrators of Apollo user accounts and roles. These G-OnRamp tools are based on tools developed by the Galaxy Genome Annotation project (https://github.com/galaxy-genome-annotation/galaxy-tools).

### 2.4 Data storage and visualization

The “CyVerse interaction via iRODS” tool facilitates the transfer of G-OnRamp output to the CyVerse Data Store for long-term storage and visualization using iRODS (https://irods.org). Genome browsers that have been produced by G-OnRamp are available through the CyVerse Data Store at https://de.cyverse.org/anon-files/iplant/home/shared/G-OnRamp_hubs/index.html.

## 3. Conclusions

G-OnRamp provides a scalable, user-friendly system for individual or collaborative eukaryotic genome annotation. G-OnRamp integrates the popular Galaxy platform, more than 25 community and custom bioinformatics tools, and the popular UCSC and JBrowse/Apollo genome browsers/annotation platforms into a powerful annotation platform. In addition to annotating novel genomes, G-OnRamp genome browsers can be used in comparative genomics studies (*e.g*., evolution of genes and pathways, understanding organism-environment interactions, etc.). G-OnRamp is useful in both research and educational settings.

